# Plant secondary metabolite-dependent plant-soil feedbacks can improve crop yield in the field

**DOI:** 10.1101/2022.11.09.515047

**Authors:** Valentin Gfeller, Jan Waelchli, Stephanie Pfister, Gabriel Deslandes-Hérold, Fabio Mascher, Gaétan Glauser, Yvo Aeby, Adrien Mestrot, Christelle A.M. Robert, Klaus Schlaeppi, Matthias Erb

**Affiliations:** Institute of Plant Sciences, University of Bern, 3013 Bern, Switzerland; Department of Environmental Sciences, University of Basel, 4056 Basel, Switzerland; Institute of Geography, University of Bern, 3012 Bern, Switzerland; Department of Plant Breeding, Agroscope, 1260 Nyon, Switzerland; Platform of Analytical Chemistry, University of Neuchâtel, 2000 Neuchâtel, Switzerland; Research contracts animals group, Agroscope, 1725 Posieux, Switzerland

**Keywords:** Plant-soil feedback, soil legacy, crop rotation, secondary metabolites, plant-microbe interactions, kernel quality, wheat, maize

## Abstract

Plant secondary metabolites that are released into the rhizosphere alter biotic and abiotic soil properties, which in turn affect the performance of other plants. How such plant-soil feedbacks affect agricultural productivity and food quality in crop rotations is unknown. Here, we assessed the impact of maize benzoxazinoids on the performance, yield and food quality of three winter wheat varieties in a two-year field experiment. Following maize cultivation, we detected benzoxazinoid-dependent chemical and microbial fingerprints in the soil. The chemical fingerprint was still visible during wheat growth, while the microbial fingerprint was no longer detected. Benzoxazinoid soil conditioning by wild-type maize led to increased wheat emergence, tillering, growth and biomass compared to soil conditioning by *bx1* mutant plants. Weed cover remained unaffected, while insect damage decreased in a subset of varieties. Wheat yield was increased by over 4% without a reduction in grain quality across varieties. This improvement was directly associated with increased germination and tillering. Taken together, our experiments demonstrate that plant secondary metabolites can increase yield via plant-soil feedbacks under agronomically realistic conditions. If this phenomenon holds across different soils and environmental conditions, optimizing plant root exudation could be a powerful, genetically tractable strategy to enhance crop yields without additional inputs.

## Introduction

Plants alter the soil they live in, and thereby modulate the growth and defense status of other plants (Bever *et al*., 1997). These so-called plant-soil feedbacks can influence plant community composition and ecosystem functions (Bennett *et al*., 2017; Teste *et al*., 2017; Mariotte *et al*., 2018). They have also been used for centuries in crop rotation schemes to reduce pest, weed and disease pressure and ultimately improve crop yields (White, 1970; van der Putten *et al*., 2013). So far however, mechanistic work on plant-soil feedbacks has rarely been applied to improve crop rotations. Thus, the benefits of this research for the engineering of crop rotations for ecological and sustainable agriculture remain limited (Mariotte *et al*., 2018).

Plant-soil feedbacks can act on a variety of plant traits. Reductions in germination via allelopathic effects of exuded chemicals for instance are common (Tawaha & Turk, 2003). Changes in defense expression leading to differences in herbivore performance and preference (Pineda *et al*., 2010; Kos *et al*., 2015; Hu *et al*., 2018b; Pineda *et al*., 2020) and alterations in susceptibility to soil pathogens have also been observed (Ma *et al*., 2017). Finally, changes in hormonal balance have been linked to effects on plant growth and biomass accumulation (Pieterse *et al*., 2014; Hu *et al*., 2018a). The response to plant-soil feedbacks is often species- and variety-specific, thus requiring detailed investigations under genetically defined conditions (van der Putten *et al*., 2013; Wagg *et al*., 2015; Hu *et al*., 2018b; Cadot *et al*., 2021a). The diversity of plant traits that can be affected call for broad phenotyping efforts that take into account ecologically and economically relevant parameters, including yield and yield quality measures of agricultural output.

Plant-soil feedbacks can act via different mechanisms, including changes in nutrient availability and chemical soil properties (Bennett & Klironomos, 2019; Schandry & Becker, 2020). Positive feedbacks in agriculture are often attributed to increased soil fertility, water retention, and improved pest control (Bennett *et al*., 2012; Tamburini *et al*., 2020). In recent years, changes in root microbial communities have received substantial attention as drivers of plant-soil feedbacks (Bever *et al*., 2012; Benitez *et al*., 2021). Various plant health benefits have been associated to the rhizosphere microbiome (Berendsen *et al*., 2012) and plant-soil feedbacks represent a promising way to harness these positive effects, including growth promotion and insect resistance in agricultural settings (Hu *et al*., 2018b; Pineda *et al*., 2020).

How do plants alter soil microbial communities? Although multiple mechanisms are likely at play, the release of small molecular weight compounds, including primary and secondary metabolites, is emerging as a major determinant of microbial community composition in the rhizosphere (Pang *et al*., 2021). Flavones, coumarins, triterpenes and benzoxazinoid secondary metabolites are known to structure the rhizosphere microbiota (Hu *et al*., 2018b; Stringlis *et al*., 2018; Huang *et al*., 2019; Voges *et al*., 2019; Yu *et al*., 2021). Flavones and benzoxazinoids have recently been shown to modulate plant-soil feedbacks via changes in microbial communities (Hu *et al*., 2018b; Yu *et al*., 2021). If and how secondary metabolites can alter plant performance via plant-soil feedbacks under realistic field conditions, however, remains unclear.

Benzoxazinoids are a class of indole-derived secondary metabolites that are produced and released in high quantities by important food crops such as maize and wheat (Frey *et al*., 2009; Hu *et al*., 2018b). Multiple functions of benzoxazinoids have been described, ranging from defense to nutrient uptake (Niemeyer, 2009). Soil conditioning by benzoxazinoids can feed back on growth and defense of maize and wheat (Hu *et al*., 2018b; Cadot *et al*., 2021a). Benzoxazinoids can shape root microbial communities (Hu *et al*., 2018b; Cotton *et al*., 2019; Kudjordjie *et al*., 2019; Cadot *et al*., 2021b), chelate iron in the soil and possibly reduce the performance of non-benzoxazinoid producing plants via allelopathic effects (Bigler *et al*., 1996; Niemeyer, 2009; Hu *et al*., 2018a). Thus, they may influence crop rotations and yields through a variety of plant-soil feedback mechanisms.

To test whether benzoxazinoids can influence crop yields in a rotation scheme, we investigated how maize benzoxazinoids affect weed and pest pressure, germination, growth, yield, and food quality of wheat plants in the field. Maize and wheat are among the most important crops in global food production and are commonly cultivated in sequence in rotation schemes. In a two-year field experiment involving wild-type and benzoxazinoid-deficient *bx*_*1*_ mutant maize plants, we first evaluated the effects of benzoxazinoid soil conditioning on soil chemistry and microbial communities. In the following season, we planted three wheat varieties into the same field and quantified a wide variety of agronomically important traits, including yield and yield quality. Through this experiment, we demonstrate that root exudation of secondary metabolites can be linked directly to improved food production under an agronomically realistic crop rotation scenario.

## Results

### Maize benzoxazinoid soil conditioning results in persistent chemical fingerprints in the soil

To test the hypothesis that maize benzoxazinoids modulate the performance of wheat in a crop rotation scheme, we grew wild-type W22 and benzoxazinoid-deficient *bx*_*1*_ mutant maize plants (in a W22 background) in the field. Compared to its wild-type counterpart, the *bx1* mutant exhibits a strong reduction in benzoxazinoid production due to a transposon insertion in the *Bx1* gene (Tzin *et al*., 2015). Previous work has shown that consistent soil conditioning and feedback effects can be triggered by different *bx1* mutant alleles in different genetic backgrounds (Hu *et al*., 2018b). The two genotypes were alternatingly sown in 5 strips, each consisting of 12 rows of maize (Fig. S1). Both maize genotypes grew similarly and accumulated the same amount of biomass at the end of the growing season, most likely due to abundant micronutrients and low pest pressure (Fig. S2A). Substantial amounts of benzoxazinoids and benzoxazinoid degradation products were detected in the soils of plots cultivated with wild-type plants (Fig. 1A). HDMBOA-Glc was the most abundant benzoxazinoid, followed by HMBOA, DIMBOA and DIMBOA-Glc. The breakdown products MBOA and AMPO were also detected. Most benzoxazinoids were below the limit of detection in the soils planted with *bx1* mutant plants. We only detected trace amounts of MBOA and AMPO in these soils.

**Figure 1.**
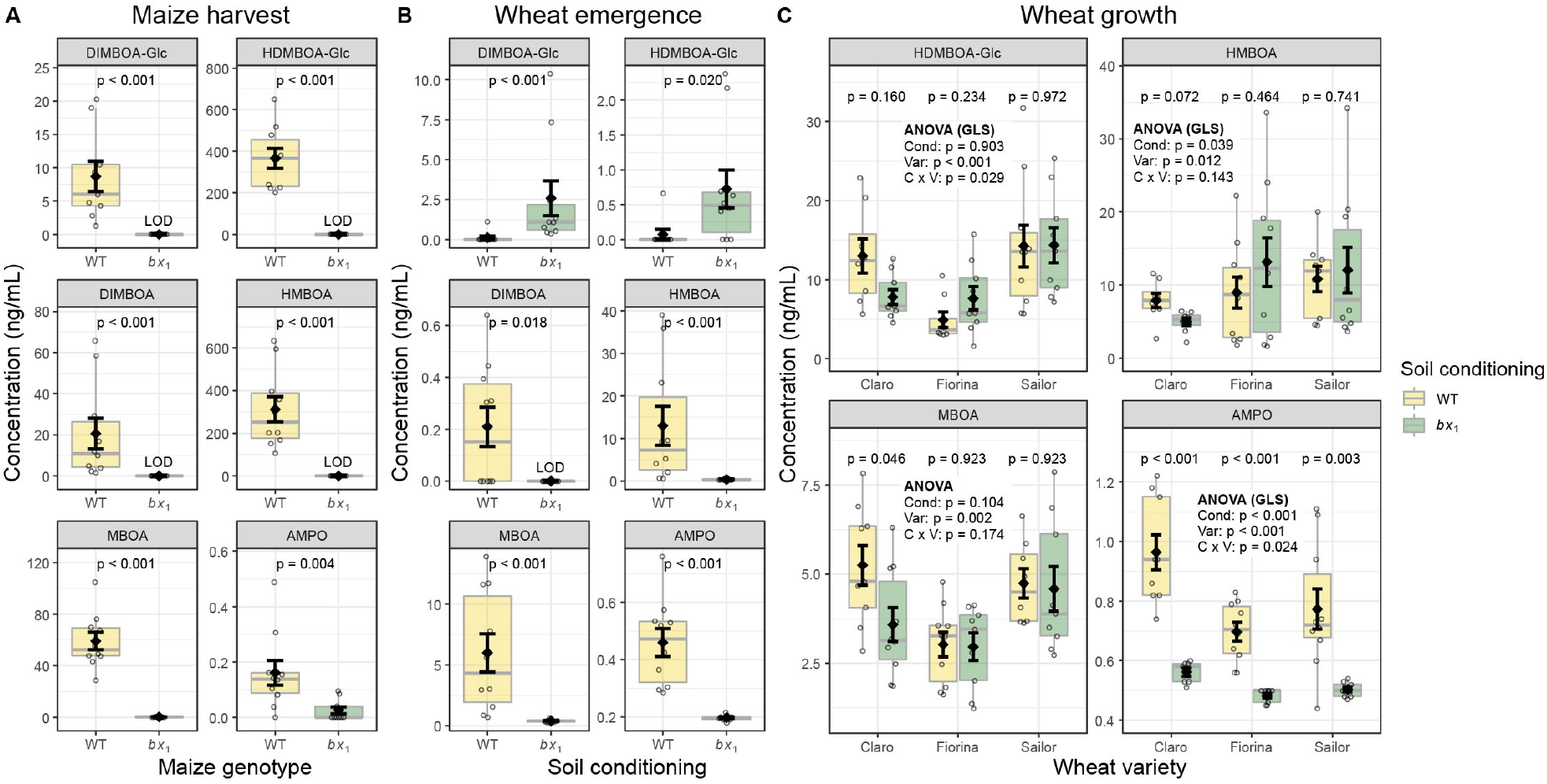
Root benzoxazinoid exudation results in persistent chemical fingerprints. **(A)** Concentrations of benzoxazinoids in soil at harvest of the wild-type (WT) or benzoxazinoid-deficient *bx*_*1*_ mutant maize plants indicated in ng per mL of soil. **(B)** Benzoxazinoids in field soil 6 weeks after maize harvest. For **(A)** and **(B)** means ± SE, boxplots, and individual datapoints are shown and Wilcoxon rank-sum tests are included (FDR-corrected *p* values, n = 10). **(C)** Benzoxazinoids were measured again during wheat growth. Means ± SE, boxplots, and individual datapoints are shown (n = 10). ANOVA tables (if unequal variance, on generalized least squares model GLS) and pairwise comparisons within each wheat variety are included (FDR-corrected *p* values). LOD: below limit of detection. Gen: maize genotype (WT and *bx*_*1*_). Cond: soil conditioning (WT and *bx*_*1*_). Var: wheat variety. ‘C x V’: interaction between conditioning and wheat variety.

To evaluate the persistence of this chemical fingerprint at the time of cultivation of the next crop, we determined benzoxazinoid profiles 6 weeks after maize harvest, at the beginning of winter wheat cultivation. Most benzoxazinoids were only present in trace amounts, with concentrations that were 3-to 800-fold lower compared to the end of maize cultivation (Fig. 1B). An exception was the stable breakdown product AMPO, which had increased more than 2-fold in abundance during this time. DIMBOA, HMBOA, MBOA and AMPO were still present in higher concentrations in wild-type conditioned soils. Interestingly, the two glycosylated benzoxazinoids DIMBOA-Glc and HDMBOA-Glc were more abundant in soils previously cultivated with *bx*_*1*_ mutant maize. As benzoxazinoids are released as glycosides and deglycosylated in the soil, this is indicative of a faster deglycosylation of benzoxazinoids in wild-type conditioned soils.

To test if soil conditioning by benzoxazinoids also affected other soil edaphic factors, we analyzed soil macro and micronutrient levels and pH at the end of maize cultivation. No significant differences were found between soils cultivated with wild-type or *bx1* mutant plants (Fig. S2B).

We then sowed 2 different wheat varieties (Claro and Fiorina) into the field, resulting in 20 plots per wheat variety with a size of 6 * 6 m where half of the plots were previously cultivated with wild-type and the other half with *bx*_*1*_ mutant maize (Fig. S1). An additional variety (Sailor) was sown for multiplication adjacent to the two other varieties on the same field by a third party. While Claro and Fiorina were managed without plant protection products, Sailor was treated with herbicides. As Sailor was sown within the premises of our conditioning experiment, we took the opportunity to also measure a subset of performance traits in this variety.

To test if the chemical fingerprint persisted further as wheat grew in soil, we analyzed the soil benzoxazinoids again during wheat growth. As benzoxazinoids are also produced by wheat, the measurements likely represent both old maize and newly wheat produced metabolites. The previous clear differences in benzoxazinoid levels of HDMBOA-Glc, HMBOA and MBOA were not detected any more at this point (Fig. 1C). However, AMPO levels were still significantly and consistently higher in plots where previously wild-type maize grew, this was apparent across all three wheat varieties. Taken together, these results show that modulating benzoxazinoid production results in a persistent soil chemical fingerprint.

### Maize benzoxazinoid soil conditioning transiently structures rhizosphere microbial communities

To investigate if differences in benzoxazinoid soil conditioning affected the bacterial and fungal communities, we analyzed soil, rhizosphere, and root samples by profiling the bacterial 16S rRNA gene and the ITS1 region of the ribosomal operon for fungi. Microbiota profiling at maize harvest revealed the biggest taxonomic differences at the phylum level to be among compartments (Fig. S3). Permutational Multivariate Analysis of Variance (PERMANOVA) on Bray-Curtis distances revealed significant differences between genotypes in the bacterial and fungal community composition of roots and rhizospheres after taking the effect of the sequencing run into account (Fig. 2A). Plant genotype accounted for 9.7 % to 15.7 % of the total variation within compartments. The benzoxazinoid effect on root bacterial and fungal community was comparable (*R*^*2*^ bacteria = 13 %, *R*^*2*^ fungi = 12.7 %) while in the rhizosphere we found a more pronounced effect on the fungal community relative to the bacterial community (bacteria = 9.7 %, fungi = 15.7 %). In bulk soil, no benzoxazinoid effects were detected (Fig. 2A). In line with PERMANOVA, visualization of bacterial and fungal communities in roots and rhizospheres by Constrained Analysis of Principal Coordinates (CAP) showed a clear differentiation between maize genotypes in both compartments (Fig. 2B). Overall, these results confirm that benzoxazinoids structure root-associated microbial communities in maize.

**Figure 2.**
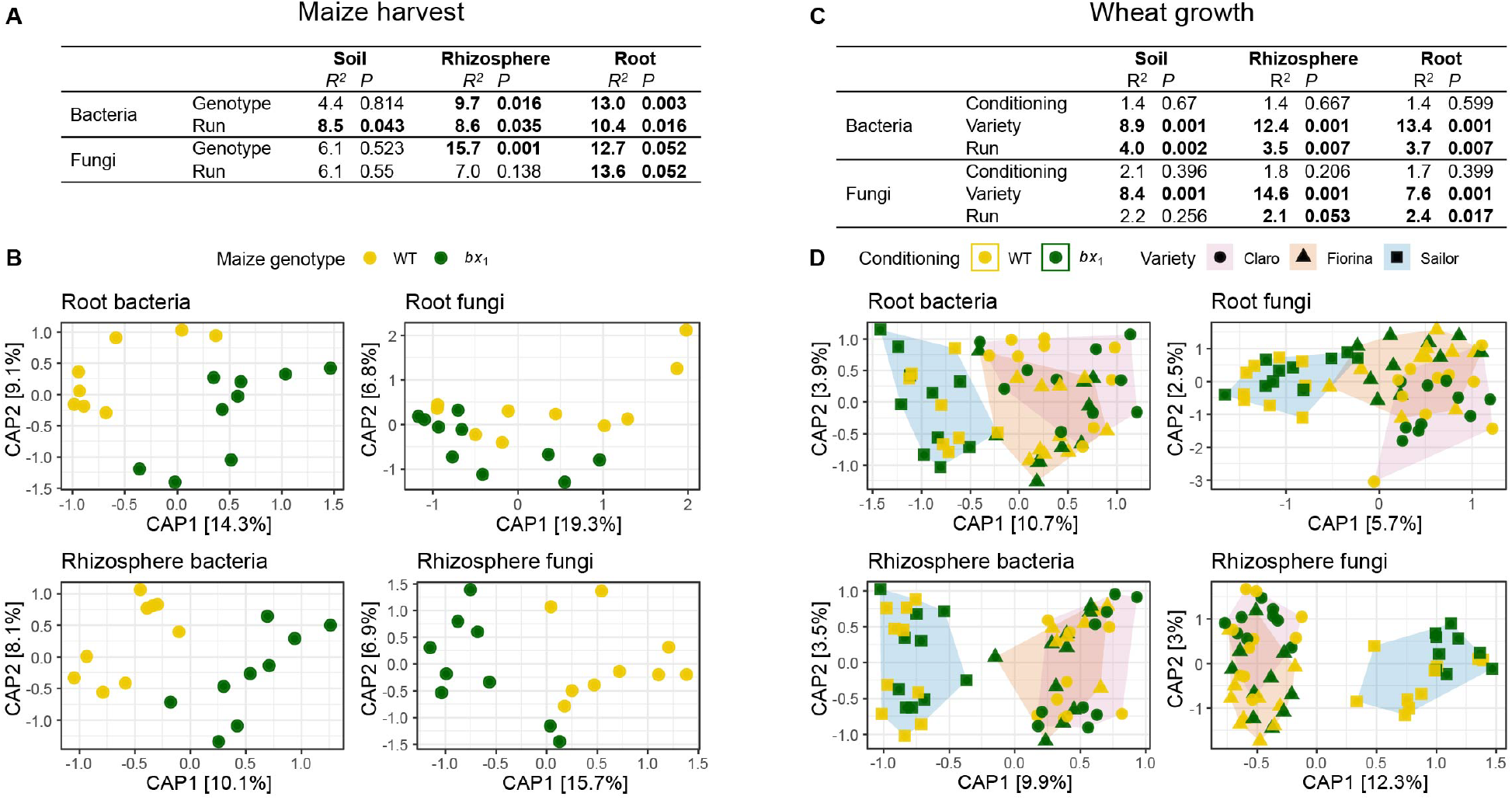
Root benzoxazinoid exudation transiently affect microbial communities. Soil, rhizosphere, and root-associated microbial communities at maize harvest (**A, B**) and during wheat growth (**C, D**). **(A)** Output of PERMANOVA on Bray-Curtis dissimilarities of bacteria and fungi showing *R*^*2*^ and *p* values for genotype and sequencing run effects in soil, rhizosphere, and root compartments. Significant effects are indicated in bold. **(B)** Constrained Analysis of Principal Coordinates (CAP) confirming the genotype effects found in the PERMANOVA, axis labels denote percentage of explained variance (n = 8-10). **(C, D)** Same as in (**A, B**) but also including the factor wheat variety (n = 6-10).

To test if the benzoxazinoid effects on microbial community composition persisted during crop rotation, we analyzed bacteria and fungi in the root, rhizosphere, and soil compartments during wheat growth. Again, the strongest taxonomic differences at phylum level were found among compartments (Fig. S4). PERMANOVA revealed a consistent difference in community composition between wheat varieties, with Sailor being the most dissimilar to the others (Fig. 2C/D). Note that these differences could also be the result of different position of Sailor in the field (Fig. S1). PERMANOVA did not reveal any benzoxazinoid-dependent effects on microbial community composition. Thus, there was no clear legacy effect on microbial community composition at the onset of wheat maturation.

### Maize benzoxazinoid soil conditioning improves subsequent wheat emergence and growth

To investigate whether benzoxazinoid soil conditioning affects wheat performance, we measured emergence shortly after seeding and leaf chlorophyll content, plant height and aboveground biomass during wheat growth of the three varieties. Overall, wheat seedling emergence was increased by 8 % in benzoxazinoid conditioned soils (Fig. 3A). Chlorophyll content measured in the youngest fully developed leaf (as a proxy for early plant performance) was generally increased in plants growing in benzoxazinoid conditioned soils (Fig. 3B). Later during wheat growth, height and biomass production per area, as well as shoot water content were increased in benzoxazinoid conditioned soils (Fig. 3C/D, Fig. S5). Thus, benzoxazinoid soil conditioning increases wheat performance across different wheat varieties.

**Figure 3.**
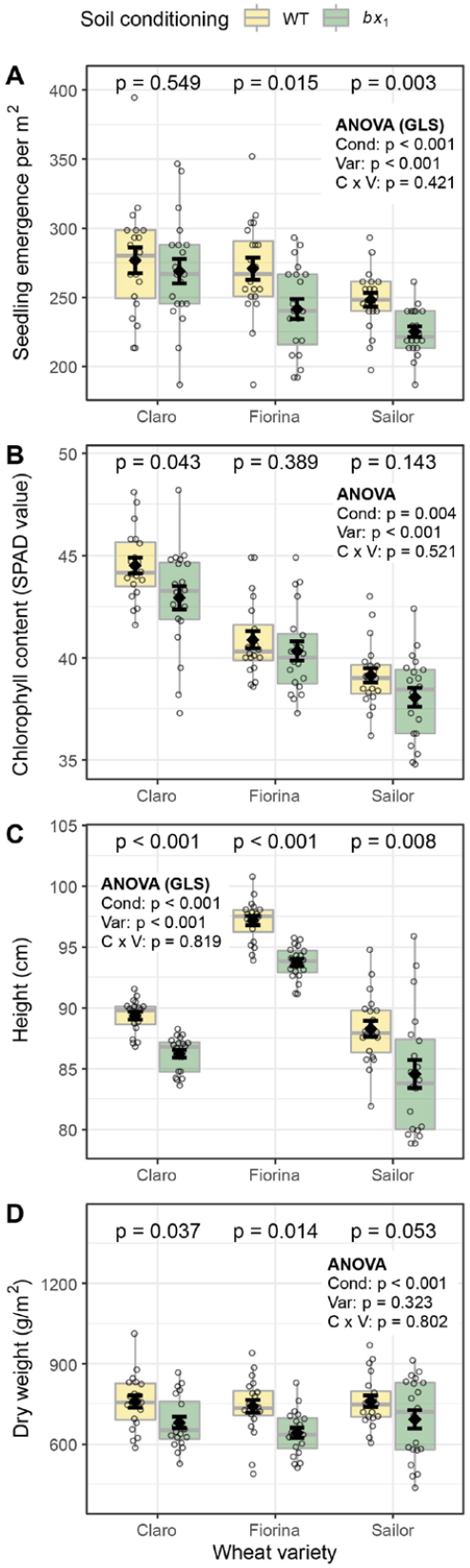
Benzoxazinoid soil conditioning positively affects wheat performance. **(A)** Seedling emergence, **(B)** chlorophyll content, **(C)** plant height, and **(D)** shoot dry weight of three wheat varieties sown in soils previously conditioned with wild-type (WT) or benzoxazinoid-deficient *bx*_*1*_ mutant maize. Means ± SE, boxplots, and individual datapoints are shown (n = 20). ANOVA tables (if unequal variance, on generalized least squares model GLS) and pairwise comparisons within each wheat variety (FDR-corrected *p* values) are included. Cond: soil conditioning (WT and *bx*_*1*_). Var: wheat variety. ‘C x V’: interaction between conditioning and wheat variety.

### Maize benzoxazinoid soil conditioning does not change weed pressure, but reduces insect infestation

To test for possible allelopathic effects of benzoxazinoids on weeds, we surveyed the weed cover on all plots. Chickweed (*Stellaria media*), Persian speedwell (*Veronica persica*), and Shepherd’s purse (*Capsella bursa-pastoris*) were the most abundant weeds on the plots. We found that weed pressure differed along the field, and therefore accounted for positional effects in the analysis. If statistically significant, we included position as a co-factor in further analyses. We found no effect of soil conditioning status on weed abundance for the varieties Claro and Fiorina (Fig. 4A). No weeds were detected with the variety Sailor, as the latter was treated with herbicides.

**Figure 4.**
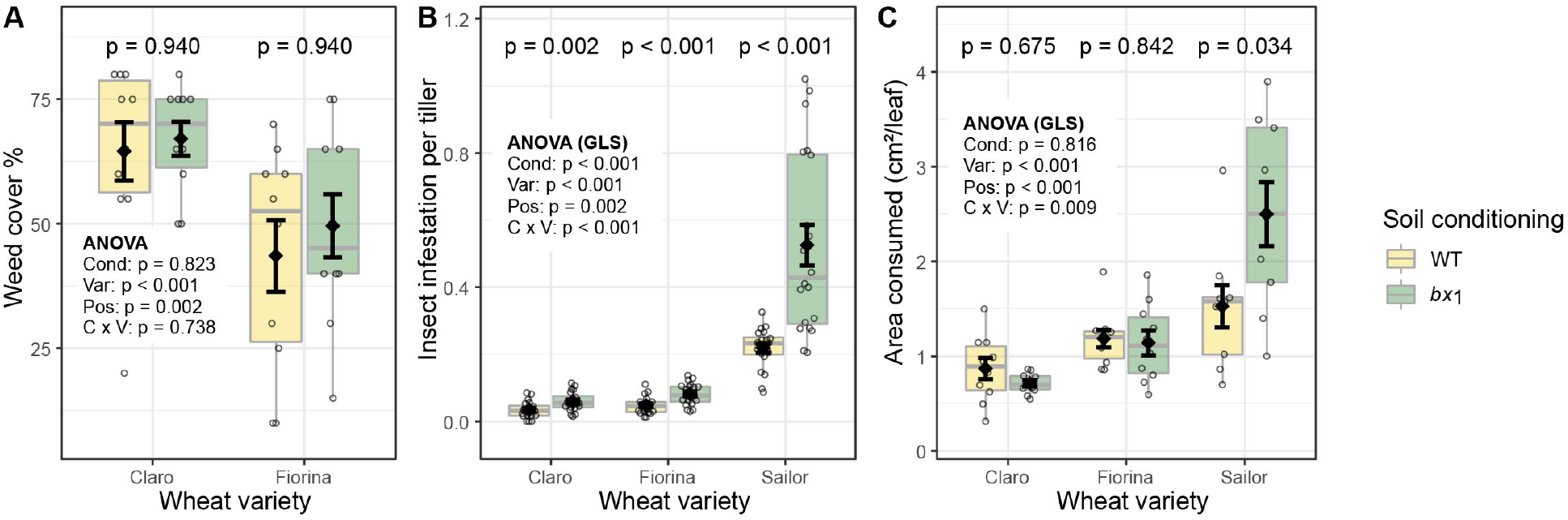
Benzoxazinoid soil conditioning reduces insect infestation on wheat, but does not affect weed pressure. **(A)** Ground cover by weed plants in plots of three wheat varieties growing in soils previously conditioned with wild-type (WT) or benzoxazinoid-deficient *bx*_*1*_ mutant maize (n = 10). No weeds were detected in plots with the variety Sailor due to herbicide treatment of this variety. **(B)** Mean abundance of cereal leaf beetle (*Oulema melanopus*) per tillers (n = 20) and **(C)** Consumed flag leaf area by cereal leaf beetle (n = 9-10). Means ± SE, boxplots and individual datapoints (n = 20) are shown. ANOVA tables (if unequal variance, on generalized least squares model GLS) and pairwise comparisons within each wheat variety (FDR-corrected *p* values) are included. Cond: soil conditioning (WT and *bx*_*1*_). Var: wheat variety. ‘C x V’: interaction between conditioning and wheat variety. Pos: position on the field.

The main herbivore that occurred in the field was the cereal leaf beetle *Oulema melanopus*. The abundance of *Oulema* larvae was significantly reduced on wheat plants of all three varieties grown in benzoxazinoid conditioned soils, with the biggest difference in the variety Sailor (Fig. 4B). To investigate whether this pattern resulted in reduced damage, we quantified the consumed leaf area on the flag leaves at the end of *Oulema* development. Benzoxazinoid conditioning reduced leaf damage in the variety Sailor, but not in Claro and Fiorina (Fig. 4C). We also measured defense hormone levels, indicative for defense activation. No significant influence of benzoxazinoid soil conditioning was found (Fig. S6).

### Maize benzoxazinoid soil conditioning increases wheat biomass upon maturation

To understand how benzoxazinoid soil conditioning influences mature wheat plants, we also quantified plant performance at harvest. All wheat varieties had a higher number of tillers per area in benzoxazinoid conditioned soils (Fig. 5A). To test if these differences can be attributed to differences in emergence or differences in tillering, we also counted the number of tillers per plant. Overall, plants in benzoxazinoid conditioned soils produced a higher numbers of tillers per plant. This pattern was consistent across all varieties, with the most pronounced difference in the variety Sailor (Fig. 5B). Next, we measured if the higher tiller density resulted in a higher aboveground biomass per area. Consistent with the results during wheat growth, benzoxazinoid soil conditioning also increased biomass at plant maturity (Fig. 5C). The weight of individual tillers was similar (Fig. 5D), demonstrating that benzoxazinoid soil conditioning increased biomass by promoting tiller density, both through enhanced germination and tillering.

**Figure 5.**
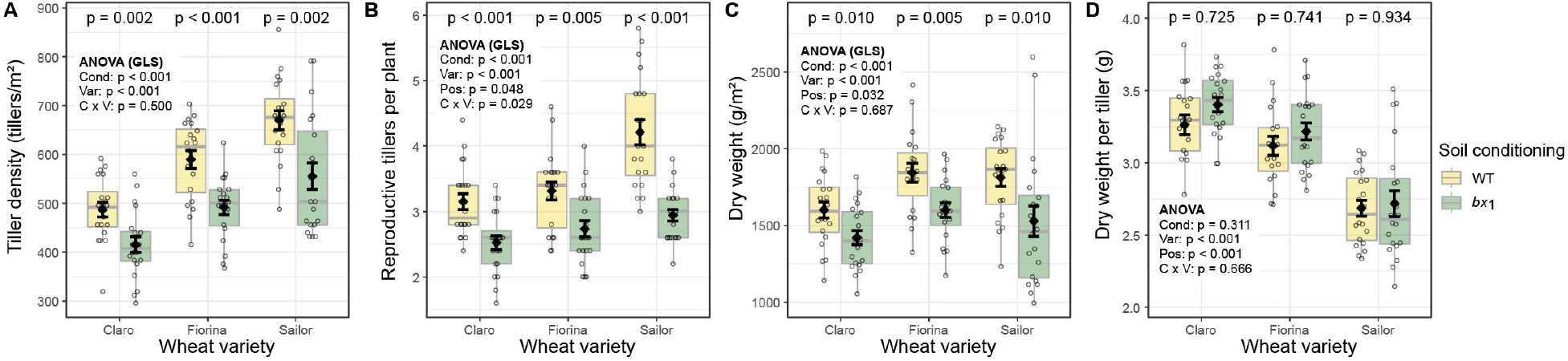
Benzoxazinoid soil conditioning increases wheat biomass in a density-dependent manner. **(A)** Tiller density, **(B)** reproductive tillers per plant, **(C)** shoot dry weight, and **(D)** dry weight per tiller of three wheat varieties growing in soils previously conditioned with wild-type (WT) or benzoxazinoid-deficient *bx*_*1*_ mutant maize. Means ± SE, boxplots, and individual datapoints (n = 20) are shown. ANOVA tables (if unequal variance, on generalized least squares model GLS) and pairwise comparisons within each wheat variety (FDR-corrected *p* values) are included. Cond: soil conditioning (WT and *bx*_*1*_). Var: wheat variety. ‘C x V’: interaction between conditioning and wheat variety. Pos: position on the field.

### Maize benzoxazinoid soil conditioning improves wheat yield

We evaluated whether benzoxazinoid soil conditioning improved wheat yield and quantified kernel weight per plot at harvest. For each Claro and Fiorina plot, 9 m^2^ were harvested. Yield was not determined for Sailor, as this field was harvested in bulk for seed multiplication. Yield in both Claro and Fiorina was increased by 4-5 % on benzoxazinoid conditioned soils (Fig. 6A). The number of kernels per tiller, the kernel weight per tiller and the thousand kernel weight did not differ between soil conditioning treatments (Fig. 6B, Fig. S7A/B), showing that the increase in yield is primarily the result of more kernels per area being produced.

**Figure 6.**
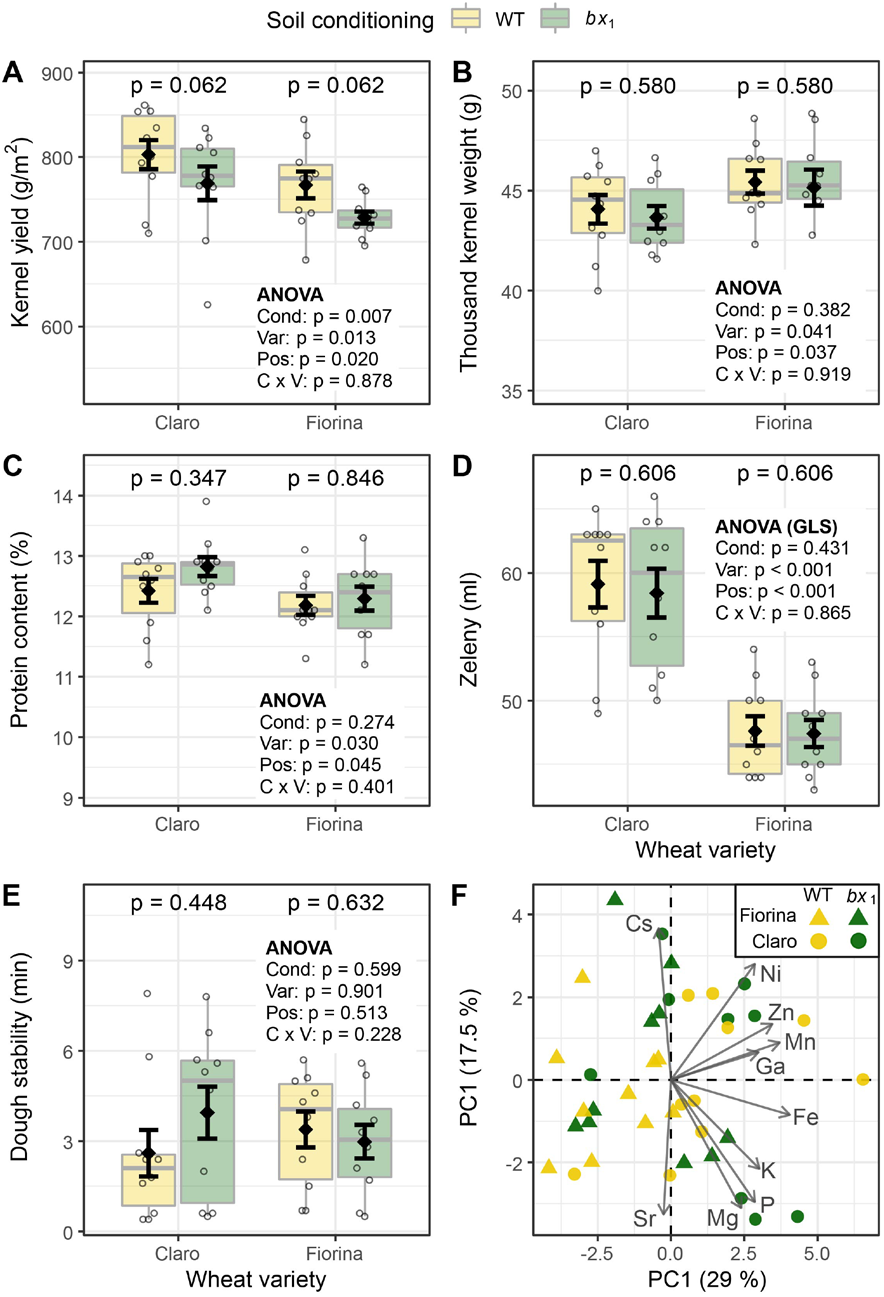
Benzoxazinoid soil conditioning enhances crop yield without a reduction in kernel quality. **(A)** Yield of two wheat varieties growing in soils previously conditioned with wild-type (WT) or benzoxazinoid-deficient *bx*_*1*_ mutant maize. Kernel quality measures included **(B)** thousand kernel weight, **(C)** kernel protein content, **(D)** Zeleny index (flour quality), **(E)** dough stability, and **(F)** PCA of kernel micronutrient composition. For **(A)-(E)** means ± SE, boxplots, and individual datapoints are shown (n = 10). ANOVA tables (if unequal variance, on generalized least squares model GLS) and pairwise comparisons within each wheat variety (FDR-corrected *p* values) are included. **(F)** reports the first two axes of the micronutrient PCA, including individual samples and the contribution of the 10 elements explaining most of the variation in the dataset (arrow length denotes relative contribution). Cond: soil conditioning (WT and *bx*_*1*_). Var: wheat variety. ‘C x V’: interaction between conditioning and wheat variety. Pos: position on the field.

To investigate whether the increased wheat yield comes with a penalty in terms of grain quality, we first determined a number of physical kernel properties. Volume per weight, kernel surface area, kernel length and kernel width were not affected by soil conditioning (Fig. S7C-F). We further assessed various agronomically important parameters that are indicative of kernel quality and suitability for baking. We measured protein content, Zeleny index, falling number, as well as dough water absorption, stability and softening. Kernel quality and baking quality were high and showed no differences between soil conditioning treatments (Fig. 6C-E, Fig. S7G-I). To test if micronutrient content is affected by soil conditioning, we also quantified 21 elements in the harvested wheat kernels. No benzoxazinoid conditioning effects were found (Fig. 6F, Fig. S8). Taken together, these results demonstrate that maize benzoxazinoid soil conditioning increases wheat yield without affecting kernel quality.

## Discussion

Plants exude secondary metabolites into the rhizosphere and thereby influence the growth and defense of subsequently growing plants (Hu *et al*., 2018b; Cadot *et al*., 2021a). Whether this phenomenon is also relevant in the field, and whether it can be exploited to improve crop productivity, is unknown. Here, we demonstrate that root secondary metabolites can improve plant growth and crop yield via plant-soil feedbacks under agronomically realistic conditions. Below, we discuss mechanisms underlying this phenomenon as well as its potential to improve sustainable food production.

Translating plant-soil feedback mechanisms to crop resistance and productivity has been proposed as a promising approach in sustainable agriculture (Mariotte *et al*., 2018). Plant secondary metabolites and their degradation products are known to suppress the growth of other plants (Schandry & Becker, 2020) and improve herbivore and pathogen resistance (Niemeyer, 2009). Less is known about their potential to influence seedling establishment (Lamichhane *et al*., 2018) and the agronomically most relevant parameters in the field, yield quality and quantity (Cadot *et al*., 2021a; Pang *et al*., 2021). We found that benzoxazinoid soil conditioning by the preceding crop increased subsequent wheat emergence, tillering and plant performance in the field, resulting in higher plant biomass and kernel yield. Because weed cover was unaffected, and increased insect infestation only resulted in increased leave damage for a subset of varieties, we conclude that the positive effects on yield were the result of directly improved germination and tillering rather than changes in plant competition or pest damage. Interestingly, the observed increase in biomass is different from what was observed in an earlier greenhouse study (2021a). This discrepancy is partially explained by the fact that the greenhouse study investigated individual plant performance and did thus not take into account germination effects. Agriculturally relevant field experiments are useful and necessary to quantify the costs and benefits of plant-soil feedbacks for sustainable agriculture.

In crop rotations the identity of the preceding crop is known to affect growth, tiller density, yield, and kernel protein content of wheat (Anderson, 2008; Rieger *et al*., 2008; Sieling & Christen, 2015). Our work expands this knowledge by demonstrating that the release of chemicals by in the preceding crop is sufficient to enhance overall crop yield through enhanced germination and tillering. Although higher plant densities are often associated with lower grain quality (Bastos *et al*., 2020), we found that the yield increase did not affect physical parameters, grain micronutrient composition, grain quality, and baking quality. The increase in yield of 4 – 5 % is equivalent to more than two years of breeding (Le Gouis *et al*., 2020), and represents a true advantage because quality remained constant without additional agricultural inputs. Benzoxazinoid exudation and responsiveness to benzoxazinoid soil conditioning are thus promising targets for future breeding efforts. Future crop rotations could be designed using varieties that are optimized for such traits. One can for instance envisage a scenario where high benzoxazinoid maize hybrids are selected specifically to precede highly responsive wheat cultivars. Future field experiments will have to evaluate how other crops respond to benzoxazinoid conditioning in the field and how generalizable the obtained results are across different years and locations. Furthermore, a better understanding of the genetic basis of benzoxazinoid exudation will be helpful to boost the release of benzoxazinoids into the soil. Such work will help to further unlock the potential of plant-soil feedbacks for the much needed sustainable intensification of agriculture (Hunter *et al*., 2017; Mariotte *et al*., 2018).

From a mechanistic perspective, plant-soil feedbacks can be triggered by different mechanisms (Bennett & Klironomos, 2019): the first plant generation changes soil chemistry (Schandry & Becker, 2020), root-associated microbiota (Bever *et al*., 2012) or their interaction, with changes in chemistry mediating changes in microbiota (Hu *et al*., 2018b; Yu *et al*., 2021). The persistence of chemical and microbiological changes is seen as a key factor in this context. It has been proposed that chemical changes may be more short lived than microbial changes, as plant secondary metabolites often degrade rapidly (Bennett & Klironomos, 2019). In line with previous studies, we found benzoxazinoids alter the composition of root-associated microbes (Hu *et al*., 2018b; Cotton *et al*., 2019; Kudjordjie *et al*., 2019; Cadot *et al*., 2021b). However, these effects disappeared by the end of the vegetative growth of the next crop. By contrast, the benzoxazinoid chemical fingerprint persisted across both cultivation periods. AMPO, a microbial degradation product with a half live of months (Macías *et al*., 2004; Niemeyer, 2009), was found in higher concentrations in benzoxazinoid conditioned soils of all three wheat varieties. Thus, we conclude that the chemical fingerprint was more long-lived that the microbial fingerprint in our study. Microbial community changes may still have contributed to plant-soil feedback effects, as many of the late phenotypes (Fig. 5, Fig. 6) were associated with initial differences in germination and tillering, where transient microbiome effects could still have been stronger. More research is needed to disentangle the relative importance of chemical and microbial fingerprints which will aid to optimize the design of agroecologically smart crop rotations.

## Conclusions

This study presents a proof of concept for the utilization of plant root exuded metabolites to increase agricultural yield without additional external inputs. This opens a new avenue to optimize plant traits in crop rotations for a more sustainable agriculture. Future studies with different varieties and crop species and in a wider range of soils and under various farming regimes will help to unravel the generalizability and applicability of using exudate-mediated plant-soil feedbacks in sustainable agriculture.

## Materials and Methods

### Plant material

The field experiment was conducted in two phases, the conditioning phase with maize (*Zea mays*) and the feedback phase with wheat (*Triticum aestivum*, Fig. S1). The wild-type maize inbred line W22 and the benzoxazinoid-deficient *bx*_*1*_ transposon knockout mutant (in W22 background, (Tzin *et al*., 2015)) were grown in the conditioning phase. During conditioning, the inbred lines were surrounded by a buffer zone of the hybrid maize variety Gottardo. In the feedback phase the wheat varieties CH Claro (referred to as Claro), Fiorina, and Sailor were grown. All three wheat varieties are commonly cultivated in Switzerland (recommended varieties by Agroscope). Claro is an obligate winter wheat, Fiorina can be cultivated as winter or spring wheat, and Sailor is a common forage winter wheat variety.

### Experimental setup

The conditioning phase indicates the first season where the field was cultivated with wild-type and *bx*_*1*_ mutant maize to condition the soil with or without benzoxazinoids. ‘Benzoxazinoid soil conditioning’ refers to the process of benzoxazinoid exudation into the surrounding soil and the resulting changes in the soil (e.g. microbial community composition). In the second season, i.e. the feedback phase, wheat was grown to survey the effects of previous benzoxazinoid soil conditioning on wheat performance. To test for genotype-specific responses, we investigated two wheat varieties Claro and Fiorina. In addition, the seed company Saatzucht Düdingen has grown a third wheat variety (Sailor) adjacent to our two wheat varieties, and we were kindly allowed to phenotype a subset of phenotypes for that variety as well. Therefore, we had three wheat varieties to survey during growth, but could not obtain data on yield and kernel quality for Sailor (Fig. S1). At the end of the conditioning phase maize biomass, belowground microbiota and soil parameters, including benzoxazinoids, were measured. In the feedback phase we determined wheat emergence, growth, and weed and insect infestation. Soil benzoxazinoids and microbiota were analyzed again during wheat growth. At the end of the feedback phase kernel quantity and quality were evaluated (Fig. S1). For detailed methods see below.

### Field specifications

The experiment was carried out in 2019 and 2020 on a field at the Agroscope research station in Posieux, Switzerland (parcel 2.3, 46°46′23.09″N 7°06′22.95″E). The soil was classified as a sandy loam. The cropping history of this field was a fodder meadow (mixture of red clover and Italian ryegrass; 2018), winter barley (2017), triticale and alfalfa (field divided, 2016), maize and alfalfa (field divided, 2015), alfalfa and maize (field divided, 2014), and alfalfa (2012-2013). The crops were managed according to Swiss conventional agricultural practices by the field team of Agroscope and the education farm of the Agricultural competence center in Grangeneuve, nearby Posieux. There was a long-lasting drought period in spring 2020 (feedback phase).

### Maize conditioning phase

Wild-type and *bx*_*1*_ inbred lines were alternately sown in 5 strips of 12 rows each (Fig. S1). Distance between maize rows was 75 cm, distance between plants within a row was 15 cm. The inbred lines were surrounded by a minimum of 18 rows of hybrid maize. Before sowing, the soil was fertilized with manure (40 m^3^/ha), ploughed, and harrowed. Weeds were once treated with herbicide (Equip Power 1.5 l/ha). During plant growth, maize was fertilized twice, firstly with ammonium nitrate supplemented with sulfur 100 kg/ha (25% N, 5% Mg, 8.5% S) and secondly with urea 180 kg/ha (46% N). Maize was harvested and silaged after 22 weeks. One week before harvest, 4 plants per maize strip were randomly selected for phenotyping resulting in 20 replicates per genotype (wild-type and *bx*_*1*_). The aboveground biomass was harvested, dried at 80 °C and weighed. For half of the samples (n = 10) soil cores of 20 × 20 × 20 cm containing the root system were excavated and used for analysis of benzoxazinoid concentrations, microbiomes, and further soil parameters as described below.

### Overview wheat feedback phase

The wheat varieties were sown one week after maize harvest. Claro and Fiorina were sown in two alternating strips, each perpendicular to the orientation of the maize rows (Fig. S1). Sailor was sown in the same orientation as the maize. Distance between wheat rows was 12.5 cm. Prior to sowing the soil was harrowed. During plant growth, wheat was fertilized twice, first with 50 kg N/ha of urea-ammonium nitrate solution (UAN; 39 % N) combined with 120 kg/ha Kiserite (15% Mg, 20% S) and second with 55 kg N/ha of UAN solution (39 % N). No plant protection products were applied to Claro and Fiorina, whereas the field of Sailor was treated with a herbicide against weeds. 4 weeks after sowing, at wheat emergence, soil samples were taken for benzoxazinoid analysis. With a soil sampler, 10 soil cores per plot (17 mm diameter, 20 cm deep)were taken and combined to one sample (n = 10 per soil conditioning). Germination, plant growth, and insect infestation were phenotyped as described below. During wheat growth, at the end of the vegetative phase, soil cores (7 × 7 cm wide, 12 cm deep) were taken below 3 randomly selected wheat plants per plot and pooled for benzoxazinoid and microbiome analysis (n = 10 per treatment combination). After 41 weeks of growth, the wheat was harvested (see below).

### Phenotyping feedback phase

To survey benzoxazinoid-dependent plant-soil feedbacks on wheat growth, we measured various parameters. Phenotyping was carried out on all subplots (Fig. S1), resulting in 20 replicates for each combination of soil conditioning status (wild-type, *bx*_*1*_) and wheat variety (Claro, Fiorina, Sailor). Weed cover estimation, determination of insect damage, and harvesting was done on plot level, resulting in 10 replicates for each treatment combination.

Emerged seedlings were counted on 1.5 m of a randomly selected wheat row within each subplot one month after the wheat was sown. Seedling emergence per m^2^ was calculated. At the end of tillering, we measured chlorophyll content with a SPADE-502 chlorophyll meter (Konica Minolta, Japan). Chlorophyll was determined in the middle of the youngest fully expanded leaf of 20 randomly selected plants per subplot and the mean value was recorded. During stem elongation, weed abundance was surveyed by estimating percentage weed cover per plot. At the end of the vegetative growth stage plant height of 10 randomly selected plants per subplot was measured and averaged for analysis. In addition, biomass accumulation was measured, by harvesting wheat plants along 1 m of a randomly selected row per subplot. Fresh biomass was weighed before plant material was dried at 80°C until constant weight, dry biomass was determined, and plant water content was calculated.

Infestation by the cereal leaf beetle (*Oulema melanopus*) was surveyed at the end of stem elongation. Along 9 m of a row within a subplot all larvae were counted and infestation per m^2^ was calculated. To determine the total larval damage on the leaves, 10 flag leaves were sampled per plot before the leaves started to wilt. Leaves were transported to the laboratory in a wettened plastic bag stored in a cooled container. Leaves were then scanned and the consumed area per leaf was determined using the R packages *EBImage* and *pliman* (Pau *et al*., 2010; Tiago Olivoto, 2021). In addition, at the end of the vegetative phase five flag leaves of five plants per plot were randomly selected, wrapped in aluminum foil and snap frozen in liquid nitrogen for later determination of phytohormone levels (see below).

Once the kernels were ripe, total biomass accumulation was determined by harvesting wheat plants along 1 m of a randomly selected row per subplot. Plant material was dried at 80 °C before measuring biomass. To calculate tiller density and weight per tiller, the number of tillers in the dried material were counted. A subsample of five randomly selected heads were threshed with a laboratory thresher (LT-15, Haldrup GmbH), and kernels were counted and weighed. Next, we randomly selected five plant per subplot and counted the number of tillers per plant, mean tiller number per plant was taken for statistical analysis.

At the end of the feedback phase, we harvested the experiment plots with a compact plot combine harvester (Zürn 110, Zürn GmbH). Yield was determined based on a 9 m^2^ area in the center of the plots (Fig. S1) and kernel weight per plot was determined. A subset of these kernels was taken for analyzing kernel quality and micronutrient composition (see below).

### Benzoxazinoid analysis

At the end of maize growth, at wheat emergence and during wheat growth soils were sampled as described above and benzoxazinoids and break down products were analyzed. Each soil sample was processed with a test sieve (5 mm mesh size), then 25 mL of soil was transferred into a 50 mL centrifuge tube and homogenized in 25 mL acidified MeOH/H2O (70:30 v/v; 0.1% formic acid). For extraction, the suspension was incubated for 30 minutes at room temperature on a rotary shaker, followed by a centrifugation step (5 min, 2000 g) to sediment the soil. The supernatant was passed through a filter paper (Grade 1; Size: 185 mm; Whatman, GE Healthcare Live Sciences), 1 mL of the flow through was transferred into a 1.5 mL centrifuge tube, centrifuged (10 min, 19000 g, 4 °C), and the supernatant was sterile filtered (Target2TM, Regenerated Cellulose Syringe Filters. Pore size: 0.45 μm; Thermo Scientific) into a HPLC glass tube for further analysis.

To obtain detectable concentrations of benzoxazinoids at wheat emergence and during wheat growth, the samples needed to be concentrated before the second centrifugation step 20 and 10 times, respectively. To obtain that, 20 mL or 10 mL of each sample was completely evaporated (45 °C; CentriVap, Labconco) and the pellet was resuspended in 1 mL of acidified MeOH/ H2O (70:30 v/v; 0.1% formic acid).

The benzoxazinoid analysis was performed as previously described (Robert *et al*., 2017). Briefly, an Acquity UHPLC system coupled to a G2-XS QTOF mass spectrometer equipped with an electrospray source and piloted by the software MassLynx 4.1 (Waters AG, Baden-Dättwil, Switzerland) was used. Gradient elution was performed on an Acquity BEH C18 column (2.1 × 50 mm i.d., 1.7 mm particle size) at 90-70% A over 3 min, 70-60% A over 1 min, 40-100% B over 1 min, holding at 100% B for 2.5 min, holding at 90% A for 1.5 minutes where A = 0.1% formic acid/water and B = 0.1% formic acid/acetonitrile. The flow rate was 0.4 mL/min. The temperature of the column was maintained at 40 °C, and the injection volume was 1 μL. The QTOF MS was operated in positive mode. The data were acquired over an m/z range of 50-1200 with scans of 0.15 seconds at a collision energy of 4 V and 0.2 seconds with a collision energy ramp from 10 to 40 V. The capillary and cone voltages were set to 2 kV and 20 V, respectively. The source temperature was maintained at 140 °C, the desolvation was 400 °C at 1000 L/hr and cone gas flow was 50 L/hr. Accurate mass measurements (< 2 ppm) were obtained by infusing a solution of leucin encephalin at 200 ng/mL and a flow rate of 10 mL/min through the Lock Spray probe. Absolute quantities were determined through standard curves of pure compounds. For that MBOA (6-methoxy-benzoxazolin-2(3H)-one) was purchased from Sigma-Aldrich Chemie GmbH (Buchs, Switzerland). DIMBOA-Glc (2-O-β-D-glucopyranosyl-2,4-dihydroxy-7-methoxy-2H-1,4-benzoxazin-3(4H)-one) and HDMBOA-Glc (2-O-β-D-glucopyranosyl-2-hydroxy-4,7-dimethoxy-2H-1,4-benzoxazin-3(4H)-one) were isolated from maize plants in our laboratory. DIMBOA (2,4-dihydroxy-7-methoxy-2H-1,4-benzoxazin-3(4H)-one), HMBOA (2-hydroxy-7-methoxy-2H-1,4-benzoxazin-3(4H)-one), and AMPO (9-methoxy-2-amino-3H-phenoxazin-3-one) were synthesized in our laboratory.

### Soil analysis

A subsample of the soil of each root system excavated at the end of maize growth (see above), was taken and pooled to obtain 4 representative samples of the field per genotype. Soil parameters were then analyzed by LBU Laboratories (Eric Schweizer AG, Thun, Switzerland). Water (H_2_O), ammonium acetate EDTA (AAE), and carbon dioxide saturated water (CO_2_) extractions were performed for different nutrients. H_2_O extracts serve as a proxy for plant available nutrients, AAE extracts for nutrients available through plant chelation mechanisms and CO_2_ extracts are a common extraction procedure for magnesium, phosphorus, and potassium (similar to H_2_O extracts).

### Phytohormones analysis

Concentrations of salicylic acid (SA), oxophytodienoic acid (OPDA), jasmonic acid (JA), jasmonic acid-isoleucine (JA-Ile) and abscicic acid (ABA) were determined by UHPLC-MS/MS. First, wheat leave samples were ground to a fine powder under constant cooling with liquid nitrogen. An aliquot of 100 mg (± 20%) was taken and the exact weight was noted for the final determination of hormone concentration. Next, phytohormones were extracted as described in Glauser, Vallat, & Balmer (2014) with minor adjustments: 10 μL of labelled internal standards (d5-JA, d6-ABA, d6-SA, and 13C6-JA-Ile, 100 ng/mL in water) were added to the samples and hormones were extracted in ethylacetate/formic acid (99.5:0.5, v/v), the samples were centrifuged and evaporated to dryness, and finally resuspended in 200 μL of MeOH 50% for analysis. Two μL of extract were injected in an Acquity UPLC (Waters, USA) coupled to a QTRAP 6500, (Sciex, USA). Analyst v.1.7.1 was used to control the instrument and for data processing. Each phytohormone peak was normalized to that of its corresponding labelled form except that of OPDA which was normalized to that of 13C6-JA-Ile.

### Kernel analysis

For morphological analysis of kernels, a subsample of 25 mL of kernels was taken. Volume weight, thousand kernel weight (TKW), kernel surface area, kernel length, and kernel width were determined by means of a microbalance and a MARVIN kernel analyzer (GTA Sensorik GmbH, Germany). A subset of kernels was milled for further analysis. To test flour quality, we determined the falling number (according to ICC standard method 107/1), Zeleny index (according to ICC standard method 116/1) and protein content, which was evaluated by near-infrared reflectance spectroscopy (NIRS) using a NIRFlex N-500 (Büchi Labortechnik AG, Switzerland). We further tested dough quality using a micro-doughLAB farinograph (model 1800, Perten Instruments, PerkinElmer United States). Dough stability (min), dough softening (Farinograph Units, FU), and water absorption capacity of the flour (%) during kneading were analyzed according to the manufacturer’s protocol.

### Kernel micronutrient analysis

We analyzed total element concentrations for 21 elements as grain micronutrients. 40 g of kernels per plot were ground to fine powder using a cutting mill (Pulverisette, Fritsch). Element extraction and analysis was performed as previously described (Cadot *et al*., 2021b), with small adjustments: An aliquot of 250 mg grain powder was extracted in 4 ml of concentrated HNO_3_ (35%) overnight and 2 mL of H_2_O_2_ (30%) was added. Samples were vortexed for 5 seconds before microwave extraction at 95°C for 30 min. Before analysis, tubes were filled to 50 mL with HNO_3_ (1%) and centrifuged (5 minutes at 2500 rpm) to remove remaining particles. Elements in the extracts were quantified with inductively coupled plasma mass spectrometry (ICP-MS, 7700x, Agilent, USA).

### Microbiota profiling

The sampling of the soil cores on the field was describe above. To prepare the soil samples, the root system was removed from the soil core, subsequently the soil was sieved through a test sieve (mesh size 5 mm). Root and rhizosphere samples were prepared as previously reported (Hu *et al*., 2018b), with minor modifications: Root segments corresponding to -5 to -10 cm below soil level were harvested and large soil particles were removed, before washing the roots twice in a 50 mL centrifuge tube with 25 mL of sterile ddH_2_0, by vigorously shaking the tube 10 times. The wash fractions were combined, centrifuged (5 minutes at 3000 g) and the resulting pellet was frozen at -80°C for further processing (rhizosphere sample). The washed roots were freeze-dried for 72 h and subsequently milled to fine powder using a Ball Mill (Retsch GmBH; 30 seconds at 30 Hz using one 1 cm steel ball).

For DNA extraction, a subsample of 200 mg soil and rhizosphere, and 20 mg of root powder was taken. DNA from all compartments were extracted using the FastDNA™ SPIN Kit for Soil (MP Biomedicals LLC, Solon, OH, USA) following the manufacturer instruction. In brief, after adding 978 μL of sodium phosphate buffer and 122 μL of MT buffer to each aliquot, the samples were homogenized with a Retsch Mixer Mill during 40 seconds at 25 Hz. Following 10 minutes of centrifugation, 250 μL of PPS was added to the supernatant. After mixing ten times by inversion, samples were centrifuged for 5 minutes. The supernatant was mixed by inversion with 1 mL of binding matrix suspension, transferred to a SPIN™ filter and then centrifuged for 1 minute. The binding matrix was washed with 500 μL of SEWS-M and a total of 3 minutes of centrifugation was performed. The matrix was air-dried for 5 minutes, and the binding matrix was resuspended with 100 μL of DNAse/Pyrogen-Free water. After incubating 5 minutes, DNA was eluted by centrifuging for 1 minute. Extraction was performed at room temperature and all centrifugation steps were done with 14000 g. After that step, the DNA was distributed into 96-well plates in a random and equal manner. The DNA concentrations were quantified with the AccuClear® Ultra High Sensitivity dsDNA quantification kit (Biotium, Fremont, CA, USA) and diluted to 2 ng μL^-1^ using a Myra Liquid Handler (Bio Molecular Systems, Upper Coomera, Australia).

For the bacterial library, a first PCR reaction was performed with the non-barcoded 16S rRNA gene primers 799-F (AACMGGATTAGATACCCKG, Chelius & Triplett, 2001) and 1193-R (ACGTCATCCCCACCTTCC, Bodenhausen *et al*., 2013). A second PCR tagged the PCR product with custom barcodes. The first PCR program consisted of an initial denaturation step of 2 minutes at 94 °C, 25 cycles of denaturation at 94 °C for 30 seconds, annealing at 55 °C for 30 seconds, elongation at 65 °C for 30 seconds, and a final elongation at 65 °C for 10 minutes. The second PCR program was similar, with the difference that the number of cycles was reduced to 10.

For the fungal library, a first PCR reaction was performed with the non-barcoded internal transcribed spacer (ITS) region primers ITS1-F (CTTGGTCATTTAGAGGAAGTAA, Gardes & Bruns, 1993) and ITS2 (GCTGCGTTCTTCATCGATGC, White *et al*., 1990). A second PCR tagged the PCR product with custom barcodes. The first PCR program consisted of an initial denaturation step of 2 minutes at 94 °C, 23 cycles of denaturation at 94 °C for 45 seconds, annealing at 50 °C for 60 seconds, elongation at 72 °C for 90 seconds, and a final elongation at 72 °C for 10 minutes. The second PCR program was similar, with the difference that the number of cycles was reduced to 7.

All PCR reactions were performed with the 5-Prime Hot Master Mix (Quantabio, QIAGEN, Beverly, MA, U.S.A.). All PCR products and pooled library were purified with CleanNGS beads (CleanNA, Waddinxveen, The Netherlands) according to manufacturer protocol with a ratio of 1:1.

All the PCR products were quantified with the AccuClear® Ultra High Sensitivity dsDNA quantification kit (Biotium, Fremont, CA, USA) and subpooled by sample type, library type and sequencing run (Table S1). Subpools were assembled using a Myra Liquid Handler by adding an equal mass of each PCR product. For the bacterial library, the rhizosphere and root subpools were purified on an agarose gel (amplicon ∼450 bp) using the NucleoSpin Gel and PCR Clean-up kit (Macherey-Nagel, Düren, Germany), whereas all other subpools were purified with CleanNGS beads. Subpools were quantified with the Qubit™ dsDNA BR kit (Invitrogen, Thermo Fisher Scientific, Waltham, MA, USA) and equally divided into two sequencing libraries (BE09 & BE10). All samples were paired-end sequenced (v3 chemistry, 300 bp paired end) on an Illumina MiSeq instrument at the NGS platform of the University of Bern.

### Bioinformatics

The raw sequencing data is available from the European Nucleotide Archive (http://www.ebi.ac.uk/ena) with the study accession PRJEB53704 and the sample IDs SAMEA110170660 (BE09) and SAMEA110170661 (BE10). Raw reads were first quality inspected with *FastQC* and demultiplexed using *cutadapt* (Andrews, 2010; Martin, 2011). The barcode-to-sample assignments are documented in the supporting Dataset S1. With *cutadapt* we also removed primer and barcode sequences from the reads (error 0.1, no indels). We utilized the *DADA2* pipeline of Callahan et al. (Callahan *et al*., 2016; R package *DADA2* v.3.10) to infer exact amplicon sequences variants (ASVs) from the sequencing reads. The raw reads were quality filtered (max. expected errors: 0; max. N’s allowed: 0), truncated to the minimal lengths (250 bp, forward read; 170 bp, reverse) and shorter and low quality reads (truncQ=2) or reads matching PhiX were discarded. The error rates were learned for the separate sequencing runs using the *DADA2* algorithm to denoise the reads and infer true sequence variants. Next, the paired forward and reverse sequences were merged by a minimal overlap of twelve identical bases, a count table was created, and chimeras were removed using the *DADA2* scripts. Finally, the taxonomy was assigned using a *DADA2* formatted versions of the *SILVA* v.132 database (Quast *et al*., 2013; Callahan, 2018) for bacteria and the FASTA general release from *UNITE* v8.3 (Abarenkov *et al*., 2021) for fungi. The bioinformatic code is available on GitHub (https://github.com/PMI-Basel/Gfeller_et_al_Posieux_field_experiment).

### Statistical analysis

All statistical analyses were conducted in R (R Core Team, 2021). Data management and visualization was performed using the *tidyverse* package collection (Wickham *et al*., 2019). Microbiota of root, rhizosphere and soil compartments were analyzed separately for maize samples (conditioning phase) and wheat samples (feedback phase). The variation between sequencing runs was taken into account in all models. We rarefied the data (100x; depth: bacteria: 8000, fungi: 1’200), because this normalization technique efficiently mitigates artifacts of different sampling depths between sample groups (Weiss *et al*., 2017). Effects on community composition were tested by permutational analysis of variance (PERMANOVA, 999 permutations) on Bray-Curtis distances in the R package *vegan* (Oksanen *et al*., 2020). For maize, we tested for differences between genotypes (model: beta diversity ∼ genotype + run), and for wheat, we tested for effects of soil conditioning and wheat variety (model: beta diversity ∼ genotype + variety + run). We visualized the beta diversity by plotting the Canonical Analysis of Principal coordinates (CAP) using the R package *phyloseq* (McMurdie & Holmes, 2013).

Plant phenotyping data was analyzed by analysis of variance (ANOVA). Statistical assumptions such as normal distribution and homoscedasticity of error variance were inspected visually from diagnostic quantile-quantile and residual plots. If unequal variance among treatment groups was observed, a model using generalized least squares (GLS, *nlme* package) was fitted, taking into account different variances for each grouping factor (Pinheiro *et al*., 2021). Possible correlations of the response variables with the position on the field were tested, and, if significant, the position on the field was factored into the model to account for otherwise unexplained variation. For linear models in the feedback phase, we tested for soil conditioning effects within each wheat variety by comparing estimated marginal means (EMMs; *emmeans* package) and reporting false discovery rate (FDR) corrected *p* values (Benjamini & Hochberg, 1995; Lenth, 2022). Wilcoxon rank-sum tests were performed to test for differences in benzoxazinoid concentrations between wild-type and *bx*_*1*_ conditioned soil in the conditioning phase and at wheat emergence; *p* values were also FDR adjusted. Maize genotype-dependent differences on soil parameters were tested by Welch’s two-sample *t*-test and *p* values were FDR adjusted. Possible differences in element profile of wheat kernels were visualized through Principal Component Analysis (PCA, *FactoMineR* package; Lê *et al*., 2008). The 10 elements explaining most of the variance in PCA-axes 1 and 2 were visualized as arrows.

All plant phenotyping and soil chemical data can be downloaded from the DRYAD repository (doi: to be inserted). All R code (microbiome and plant phenotyping) is available on GitHub (https://github.com/PMI-Basel/Gfeller_et_al_Posieux_field_experiment).

## Acknowledgement

We thank Jean-François Rauber, Wolfram Schuwey and Raphaël Grandgirard (Agricultural competence center canton Freiburg, Grangeneuve, Switzerland) for their field assistance and Lilia Levy, Lydia Michaud and Noemi Schaad (Agroscope, Changins, Switzerland) for their help during wheat harvest. Further, we are grateful to Florian Enz and Sophie Gulliver for field and laboratory assistance. This work was supported by the Interfaculty Research Collaboration “One Health” of the University of Bern.

## Author contributions

V.G., M.E. and K.S. designed research; V.G., S.P., G.D-H., F.M., G.G. and Y.A. performed research; C.A.M.R., G.G., A.M. and M.E. contributed new reagents/analytical tools; V.G., J.W., K.S. and M.E. analyzed data; and V.G., K.S. and M.E. wrote the first draft of the paper. All authors revised the paper.

## Supplementary Materials

**Fig. S1.**
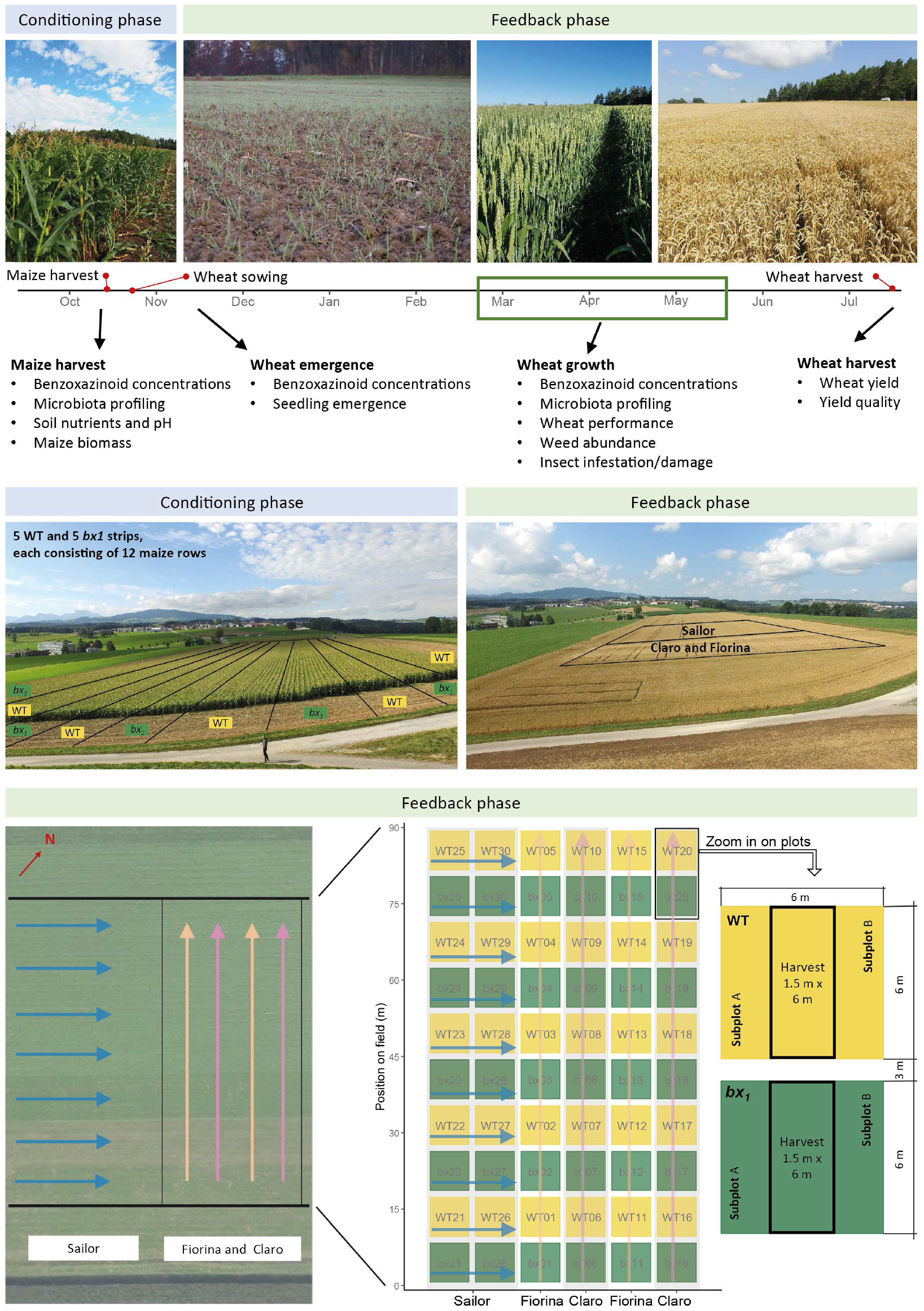
Experimental set-up. A two-year field experiment was conducted in Posieux, Switzerland. In the first season (conditioning phase) the field was cultivated with 10 strips of maize where wild-type (WT, n = 5) and the benzoxazinoid-deficient *bx*_*1*_ mutant (n = 5) maize were sown alternatingly. Each strip consisted of 12 rows of maize plants. Soil conditioning refers to the process of root benzoxazinoid exudation and the resulting changes in the soil. The impact of soil conditioning was evaluated by analysis of microbiomes, soil benzoxazinoid concentrations, soil nutrients, and pH. In the second season (feedback phase), after the maize was harvested, three wheat varieties (Claro, Fiorina, and Sailor) were sown. Most wheat phenotypes were measured in all subplots, indicated in the zoomed plots. Feedbacks of benzoxazinoid soil conditioning on wheat seedling emergence, growth, and defense were surveyed. Microbial and chemical (benzoxazinoid) soil legacies were again analyzed during wheat growth. At wheat harvest, wheat yield and yield quality were determined. For more details see the method description.

**Fig. S2.**
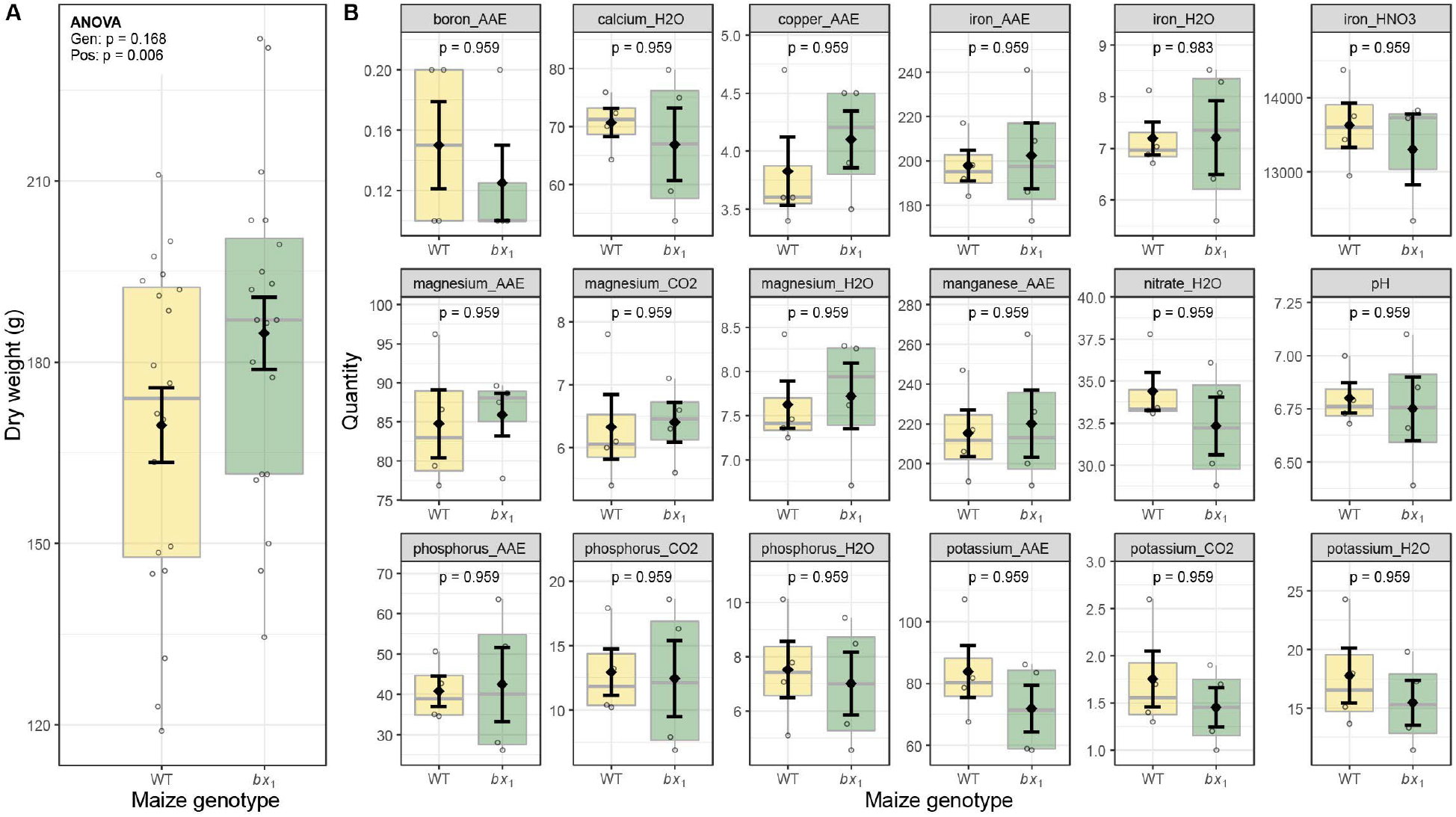
Additional parameters maize harvest. (**A**) Shoot dry weight of individual wild-type (WT) and *bx*_*1*_ maize plants at harvest. Means ± SE, boxplots, and individual datapoints are shown (n = 20). ANOVA table is included. (**B**) Soil nutrient levels at the end of the maize conditioning phase. Except for pH all values are concentrations in mg/kg soil. Means ± SE, boxplots, and individual datapoints are shown (n = 4). Welch’s two-sample *t*-tests are included (FDR-corrected *p* values). Gen: maize genotype (WT and *bx*_*1*_). Var: wheat variety. Pos: position on the field. AAE: ammonium acetate EDTA extraction; H_2_O: water extraction; CO_2_: carbon dioxide saturated H_2_O extraction.

**Fig. S3.**
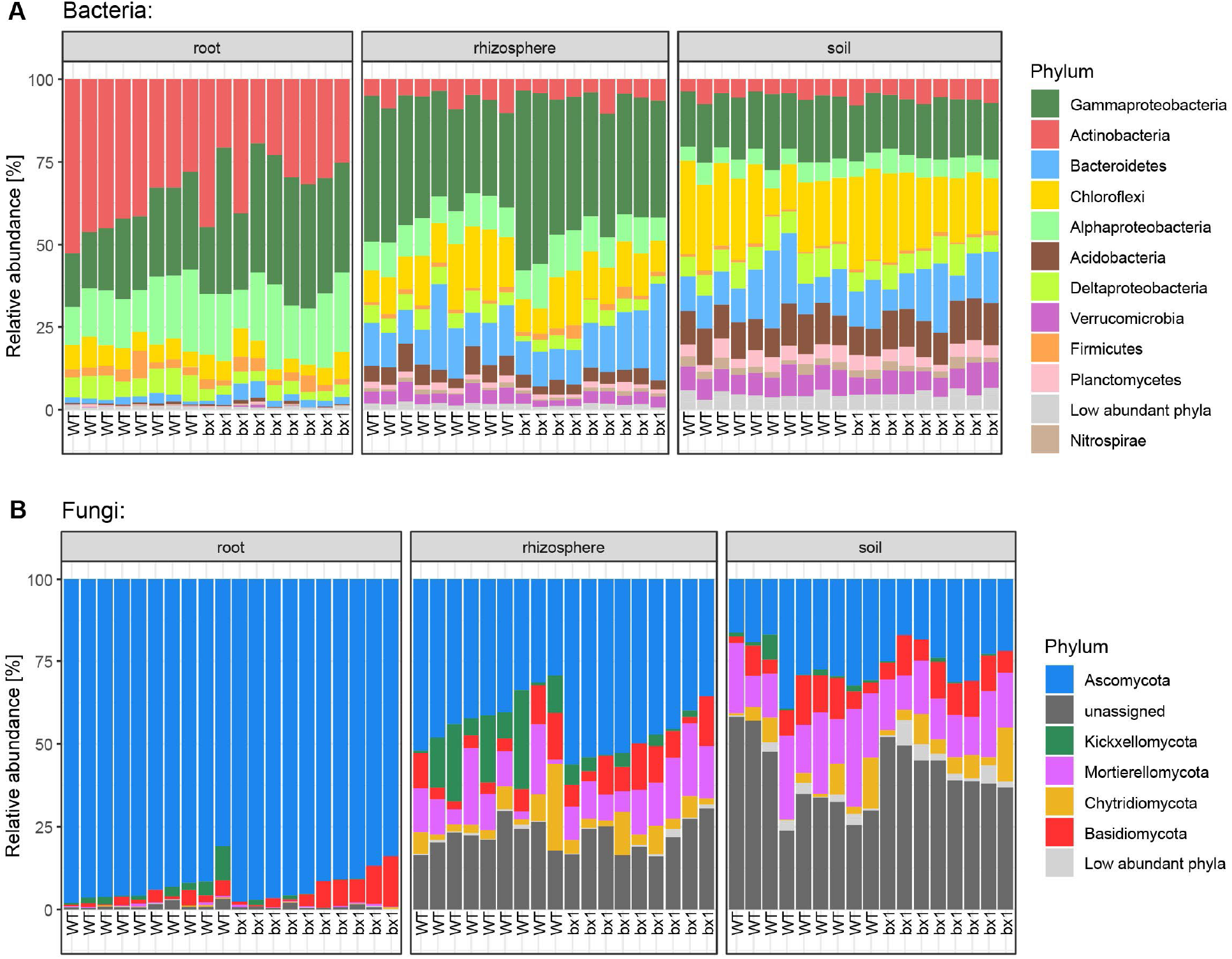
Relative abundance of microbial phyla at maize harvest. (**A**) Taxonomy of bacteria and (**B**) fungi in roots, rhizospheres, and soil of wild-type (WT) or benzoxazinoid-deficient *bx*_*1*_ mutant maize plants. All samples are shown.

**Fig. S4.**
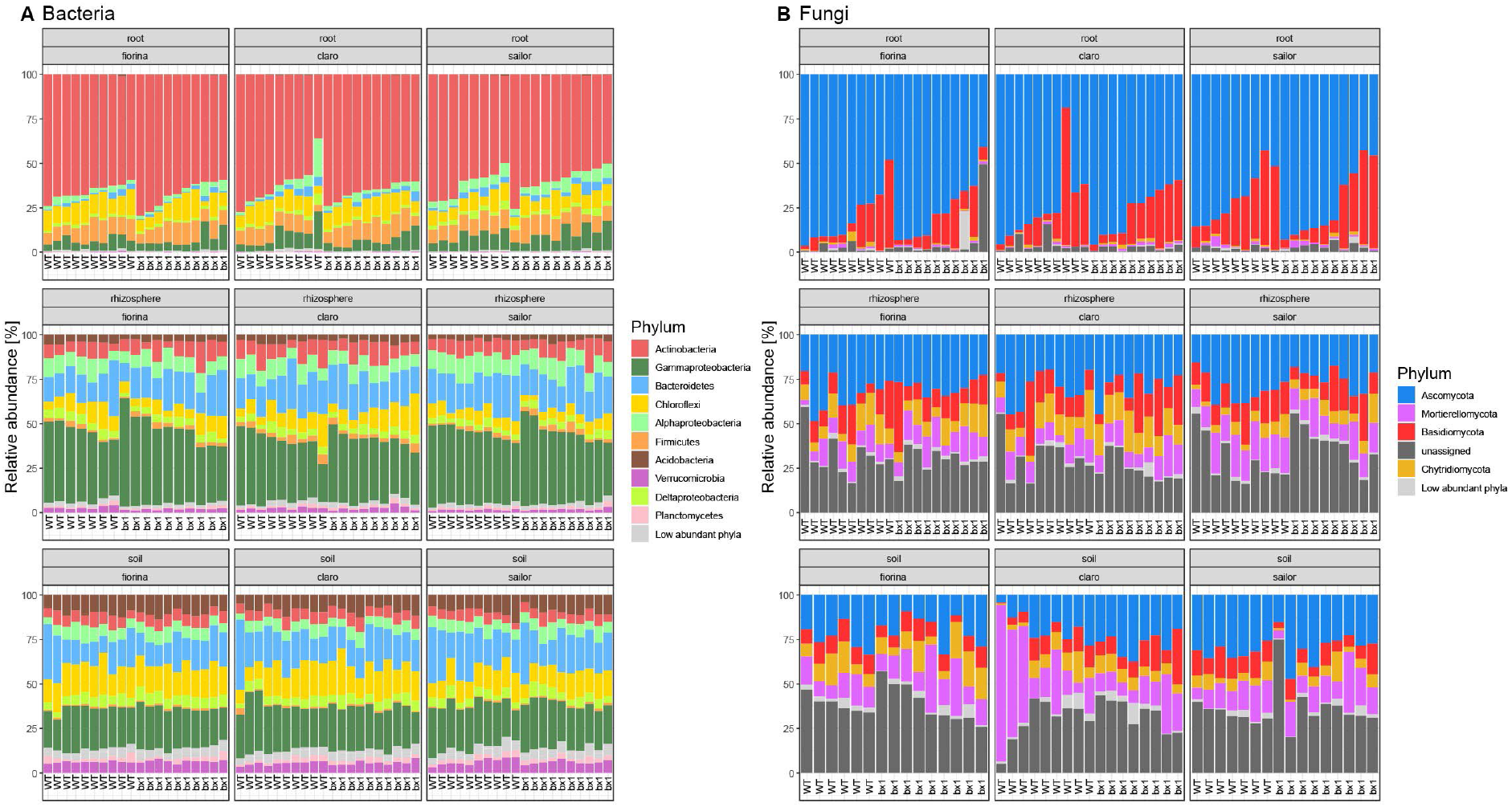
Relative abundance of microbial phyla in the wheat feedback phase. (**A**) Taxonomy of bacteria and (**B**) fungi in roots, rhizospheres, and soils in three wheat varieties grown on wild-type (WT) or benzoxazinoid-deficient *bx*_*1*_ mutant conditioned soil. All samples are shown.

**Fig. S5.**
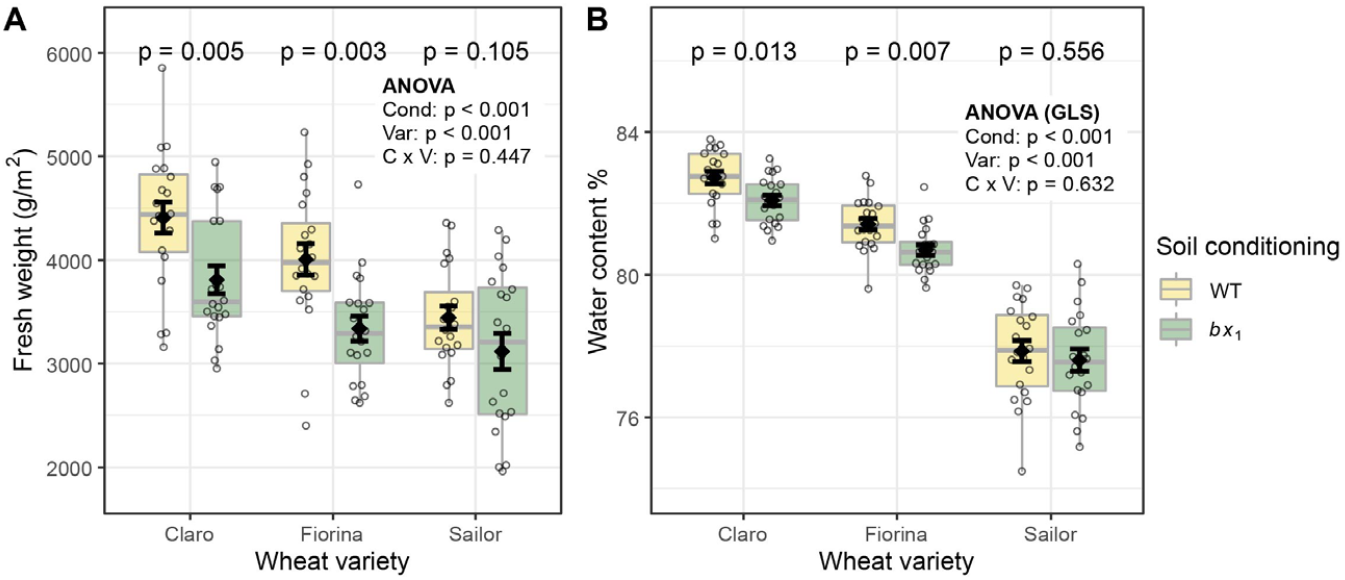
Additional benzoxazinoid soil conditioning effects on wheat growth. (**A**) Shoot fresh weight of three wheat varieties growing in soils previously conditioned with wild-type (WT) or benzoxazinoid-deficient *bx*_*1*_ mutant maize. **(B)** Shoot water content. Means ± SE, boxplots, and individual datapoints are shown (n = 20). ANOVA tables (if unequal variance, on generalized least squares model GLS) and pairwise comparisons within each wheat variety (FDR-corrected *p* values) are included. Cond: soil conditioning (WT and *bx*_*1*_). Var: wheat variety. ‘C x V’: interaction between conditioning and wheat variety.

**Fig. S6.**
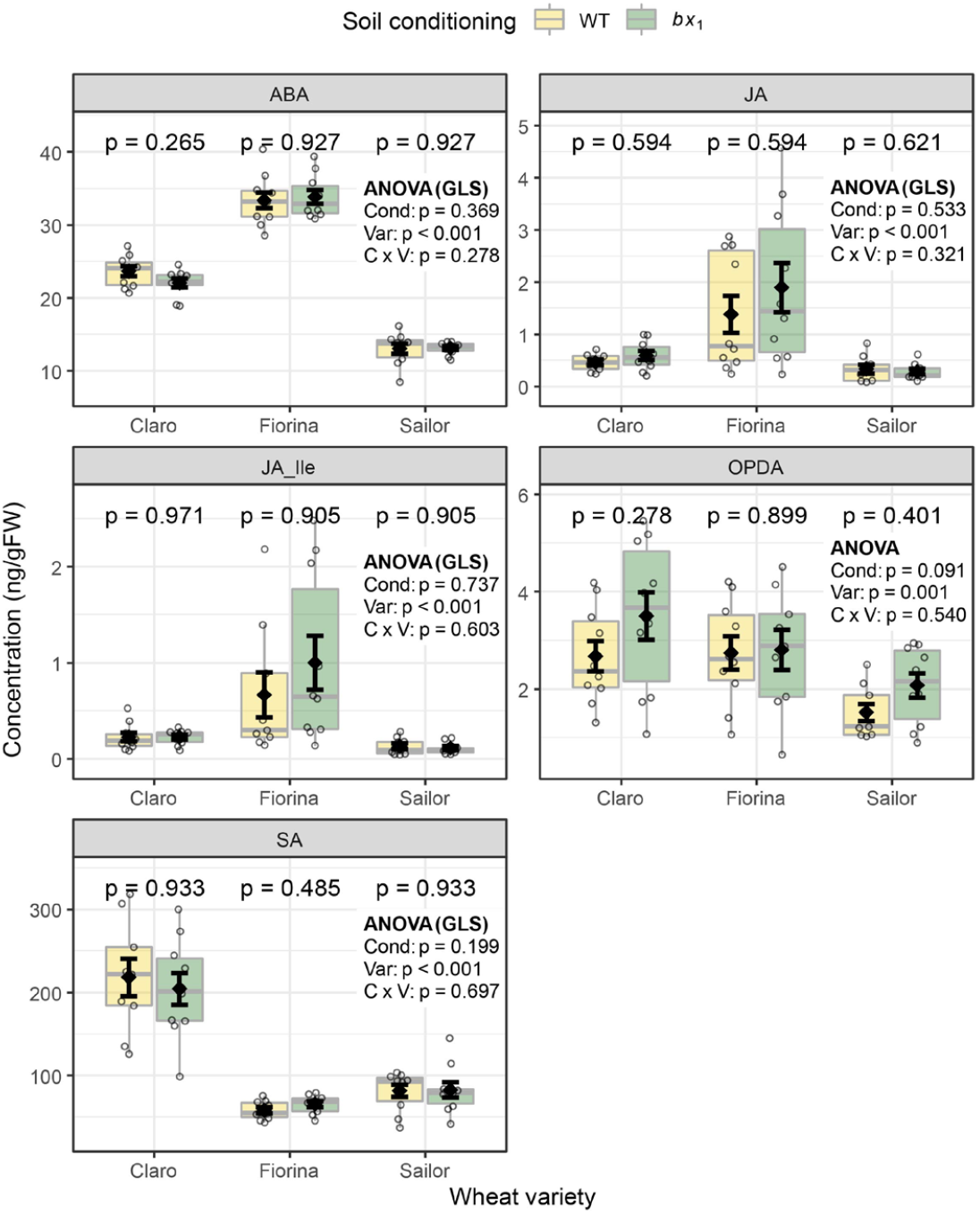
Benzoxazinoid soil conditioning does not affect leaf phytohormone levels of wheat. Phytohormone levels of three wheat varieties growing in soils previously conditioned with wild-type (WT) or benzoxazinoid-deficient *bx*_*1*_ mutant maize were measured (n = 9-10). Means ± SE, boxplots, and individual datapoints are shown. ANOVA tables (if unequal variance, on generalized least squares model GLS) and pairwise comparisons within each wheat variety (FDR-corrected *p* values) are included. FW: fresh weight. Cond: soil conditioning (WT and *bx*_*1*_). Var: wheat variety. ‘C x V’: interaction between conditioning and wheat variety. Pos: position on the field. ABA: abscicic acid. JA: jasmonic acid. JA-Ile: jasmonic acid-isoleucine. OPDA: oxophytodienoic acid. SA: salicylic acid.

**Fig. S7.**
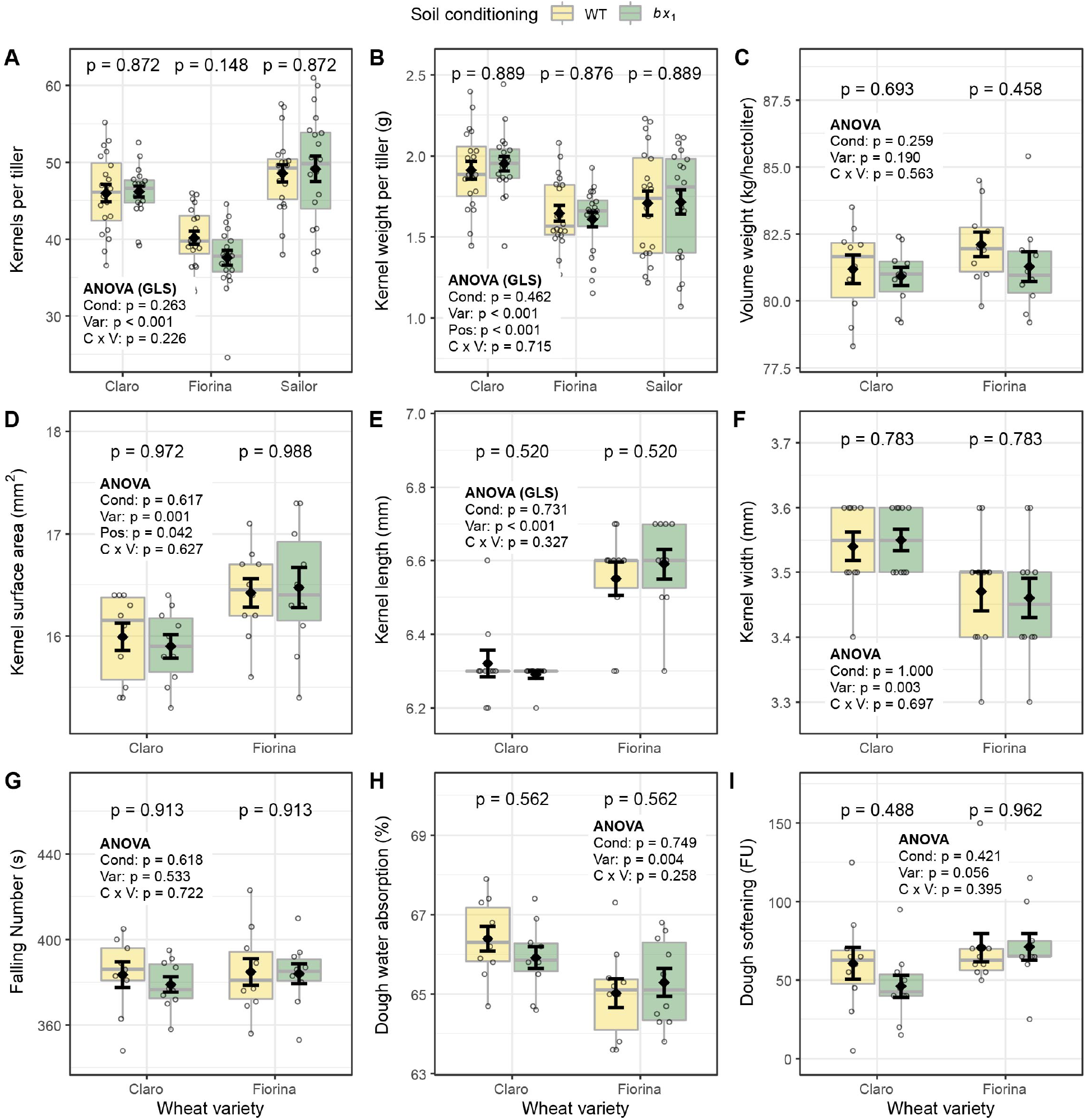
Benzoxazinoid soil conditioning does not affect wheat kernel measurements and baking quality. (**A**) Kernels per tiller of three wheat varieties growing in soils previously conditioned with wild-type (WT) or benzoxazinoid-deficient *bx*_*1*_ mutant maize (n = 10). (**B**) Kernel weight per tiller (n = 10), (**C**) kernel volume per weight (n = 10), **(D)** kernel surface area (n = 10), **(E)** kernel length (n = 10), **(F)** kernel width (n = 10), **(G)** falling number (flour quality, n = 10), **(H)** flour water absorption (n = 10), and (**I**) dough softening (n = 9-10). Means ± SE, boxplots, and individual datapoints are shown. ANOVA tables (if unequal variance, on generalized least squares model GLS) and pairwise comparisons within each wheat variety (FDR-corrected *p* values) are included. Cond: soil conditioning (WT and *bx*_*1*_). Var: wheat variety. ‘C x V’: interaction between conditioning and wheat variety. Pos: position on the field.

**Fig. S8.**
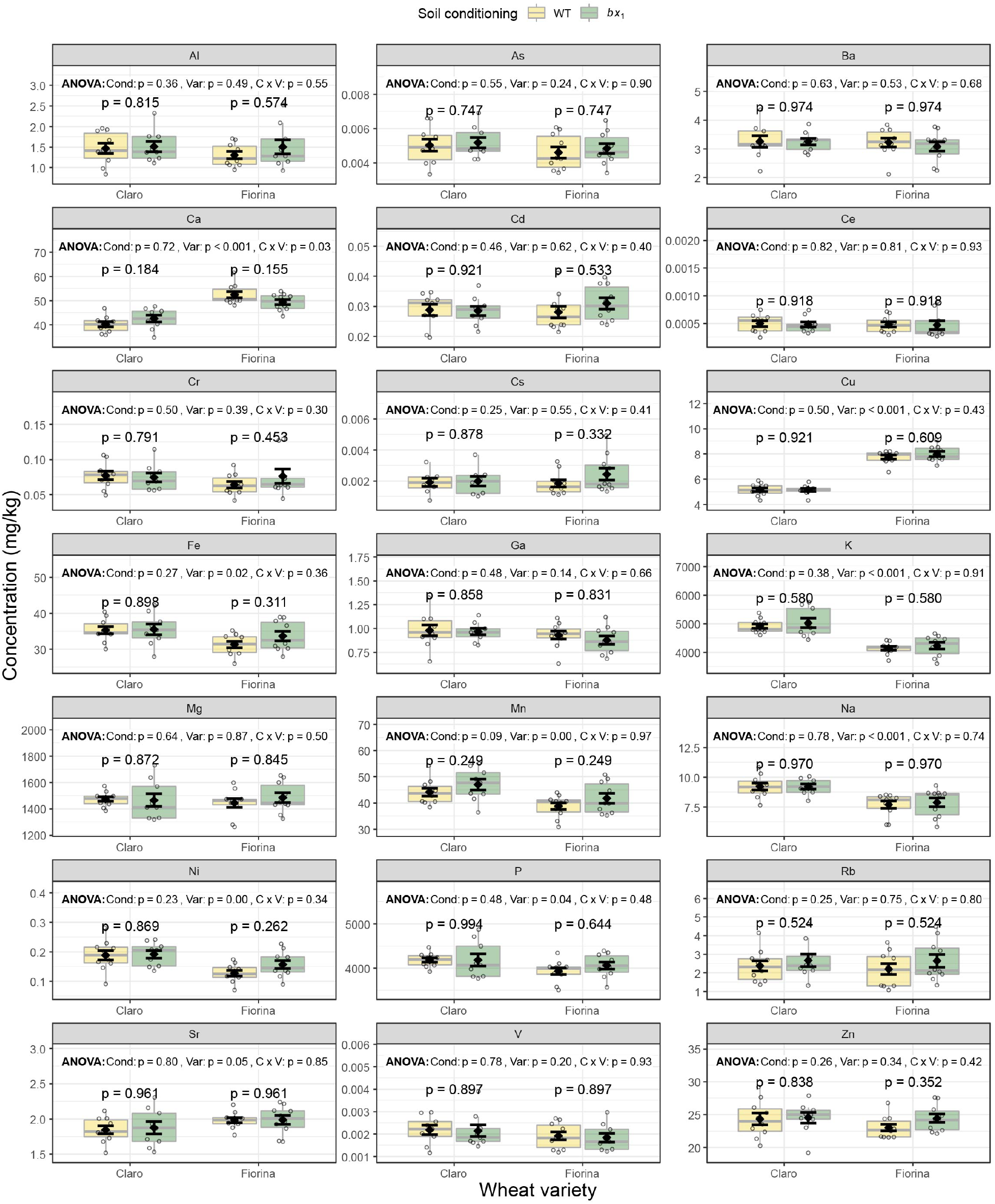
Benzoxazinoid soil conditioning does not affect micronutrient concentrations in wheat kernels. Concentration of individual elements in kernels of two wheat varieties growing in soils previously conditioned with wild-type (WT) or benzoxazinoid-deficient *bx*_*1*_ mutant maize (n = 8-10). Means ± SE, boxplots, and individual datapoints are shown. ANOVA tables and pairwise comparisons within each wheat variety (FDR-corrected *p* values) are included. Same data as shown in PCA Fig. 6F. Cond: soil conditioning (WT and *bx*_*1*_). Var: wheat variety. ‘C x V’: interaction between conditioning and wheat variety.

**Table S1.** Specifications to library preparation for sequencing.

| Sample type | Library type | Sequencing Run | Number of sample | Purification method |
| --- | --- | --- | --- | --- |
| Soil + Control | 16S (Bacteria) | BE09 | 43 | Beads |
| rhizosphere | 16S (Bacteria) | BE09 | 38 | Agarose gel |
| Root | 16S (Bacteria) | BE09 | 40 | Agarose gel |
| Soil + Control | 16S (Bacteria) | BE10 | 47 | Beads |
| rhizosphere | 16S (Bacteria) | BE10 | 42 | Agarose gel |
| Root | 16S (Bacteria) | BE10 | 40 | Agarose gel |
| Soil + Control | ITS (Fungi) | BE09 | 43 | Beads |
| rhizosphere | ITS (Fungi) | BE09 | 38 | Beads |
| Root | ITS (Fungi) | BE09 | 40 | Beads |
| Soil + Control | ITS (Fungi) | BE10 | 47 | Beads |
| rhizosphere | ITS (Fungi) | BE10 | 42 | Beads |
| Root | ITS (Fungi) | BE10 | 40 | Beads |

